# Spatiotemporal mapping of endothelial proliferative response in acute chemical liver injury

**DOI:** 10.64898/2026.10.29.760435

**Authors:** AS Rajput, H Yoon, KL Travis, EJ Hernandez, EP Shah, DB Azizoglu

**Author notes:** **Corresponding author:** D. Berfin Azizoglu (eRA Commons ID: BAZIZO). Co-first authors (AS Rajput, H Yoon).

## Abstract

Endothelial cells are major contributors to liver injury response, regeneration, and chronic liver disease. Previous studies identified a proliferative endothelial response to acute liver injury; however, a comprehensive characterization of this adaptive response is missing. We have developed a novel pipeline for spatiotemporal analysis of proliferative response in the liver. We demonstrate that endothelial cells enter the cell cycle upon acute centrilobular injury with a delay compared to hepatocytes and in a spatially distinct pattern from hepatocytes. We show that endothelial cycling predominantly occurs in the pericentral zone, while hepatocyte cycling occurs uniformly throughout the lobule. Endothelial cycling in injured livers leads to heightened clonal expansion, and to serial division rarely observed in homeostatic livers. Consequently, injured lobules exhibit endothelial overabundance during regeneration. Our study provides an in-depth characterization of endothelial proliferative response to liver injury and establishes a novel pipeline for spatiotemporal mapping of hepatic cell states.

## Introduction

Vascular endothelial maladaptation is a hallmark of chronic liver disease that affects approximately 1.5 billion people worldwide^1^. Recent clinical studies suggest that preventing endothelial maladaptation can alleviate fibrosis in chronic liver disease patients, warranting further investigation into the mechanisms of endothelial maladaptation and its reversal^2^. Hepatic endothelial maladaptation is characterized by capillarization and hypertension, and can develop because of persistent injury to the liver^3,4^. While this is a long and complex process, it initiates with acute injury. Acute toxicant or medication exposure initializes a reversible and transient adaptive response in the hepatic endothelium^5,6^. This is followed by a highly effective regenerative process where the liver fully recovers its function^5,6^. The acute liver injury setting provides an opportunity to define the mechanisms of early-onset injury response, adaptation, and the reversal of maladaptive alterations in the hepatic endothelium.

Liver regeneration following acute injury occurs within anatomical liver units termed lobules organized around landmark blood vessels^7^, 1) portal veins at the boundary, 2) a central vein in the middle, and 3) sinusoids connecting the former two vessel sets. In acute centrilobular chemical liver injury, medication or hepatotoxins such as carbon tetrachloride (CCl_4_) target hepatocytes in the pericentral zone of the lobule, causing hepatocyte necrosis^8^. The remaining hepatocytes or a subset thereof enter the cell cycle and expand to repopulate the lobule^9,10^. Over the past decade, extensive work focused on defining the proliferative hepatocyte populations and their location within the lobule^11–21^. These studies have provided evidence for enhanced cycling and expansion in the perinecrotic region^20,21^, the periportal zone^17^, the midlobular and periportal zones^16,18,19^, or uniformly throughout the lobule^12–14^. Similar to hepatocytes, endothelial cells have been reported to enter the cell cycle in response to CCl_4_ injury^22^. The analysis of endothelial cycling was limited to the sinusoidal subset and lacked spatial contextualization within the lobule^22^. Furthermore, while endothelial cycling has been observed, whether cycling leads to endothelial cell division and expansion in acutely injured livers has remained unknown. Previous findings motivate the entry hypothesis that endothelial cells enter the cell cycle in response to nearby hepatocyte necrosis and to support hepatocyte proliferation. This hypothesis can begin to be probed by determining the spatiotemporal correlation between endothelial cycling, hepatocyte necrosis, and hepatocyte proliferation.

The gaps in our understanding of the endothelial proliferative response, and the lack of a unified spatial model for hepatocyte proliferation, are in part due to the hitherto lack of a definitive lobule marker in mice. Recent work from our lab has established a definitive lobule marker in mice^23,24^. This provides a unique opportunity to revisit the spatial characteristics of hepatic cell cycling, and to determine how endothelial proliferative response impacts the lobule composition in acute liver injury.

Here, we utilize the recently identified definitive lobule marker EPCR^23,24^ combined with a novel image analysis pipeline and a genetic approach targeting the endothelium to characterize the endothelial proliferative response to CCl_4_-mediated liver injury in mice. We first develop an automated image analysis pipeline to spatially map cell type-specific cycling in healthy and injured liver lobules. Using this pipeline and high throughput analyses, we demonstrate that endothelial cells enter the cell cycle upon injury most prominently in the pericentral and midlobular zones. Parallel mapping of cycling in hepatocytes reveals lack of genuine spatial bias. The distribution of cycling for both cell types gradually returns to homeostatic patterns over the one week-long proliferative period. Genetic sparse clonal mapping within the endothelium reveals that fifty percent more endothelial cells divide and expand in response to injury compared to homeostasis. Unexpectedly, we identify serial endothelial division as a distinguishing characteristic of injured livers. Following the proliferative period, in line with endothelial cell expansion, the lobule and specifically the pericentral zone become overpopulated with endothelial cells. Our results suggest that hepatic endothelial cells respond to acute chemical injury through heightened proliferation and serial expansion resulting in endothelial overabundance in regenerating liver lobules.

## Results

### Novel image analysis pipeline enables accurate spatiotemporal mapping of cycling cells in liver lobules

Spatial mapping of hepatic cells in rodents has historically been challenging in part due to the lack of definitive lobule markers. Taking advantage of our recent discovery of EPCR as a lobule boundary marker^23^, we devised a workflow that enables reproducible spatiotemporal analysis of cycling cells in liver lobules. We outline this workflow in Figure 1, illustrated in endothelial cells. To temporally map cycling hepatic endothelial cells, we immunolabelled livers for ERG that marks all endothelial nuclei and Ki67 that marks cycling nuclei. We used EPCR to identify lobules. We then fed individual lobule images to the analysis pipeline. The pipeline utilizes CellProfiler^25^ followed by an in-house generated MATLAB script: First, CellProfiler is used to outline ERG ^+^ and Ki67 ^+^ cells in a lobule using custom thresholding and filters (Figure 1b-c’’). Second, a mask of the ERG^+^Ki67^+^ colocalization of each lobule is generated as an output (Figure 1d-d’’). Next, the percentage of cycling endothelial nuclei out of all endothelial nuclei is calculated across time points for temporal mapping.

**Figure 1.**
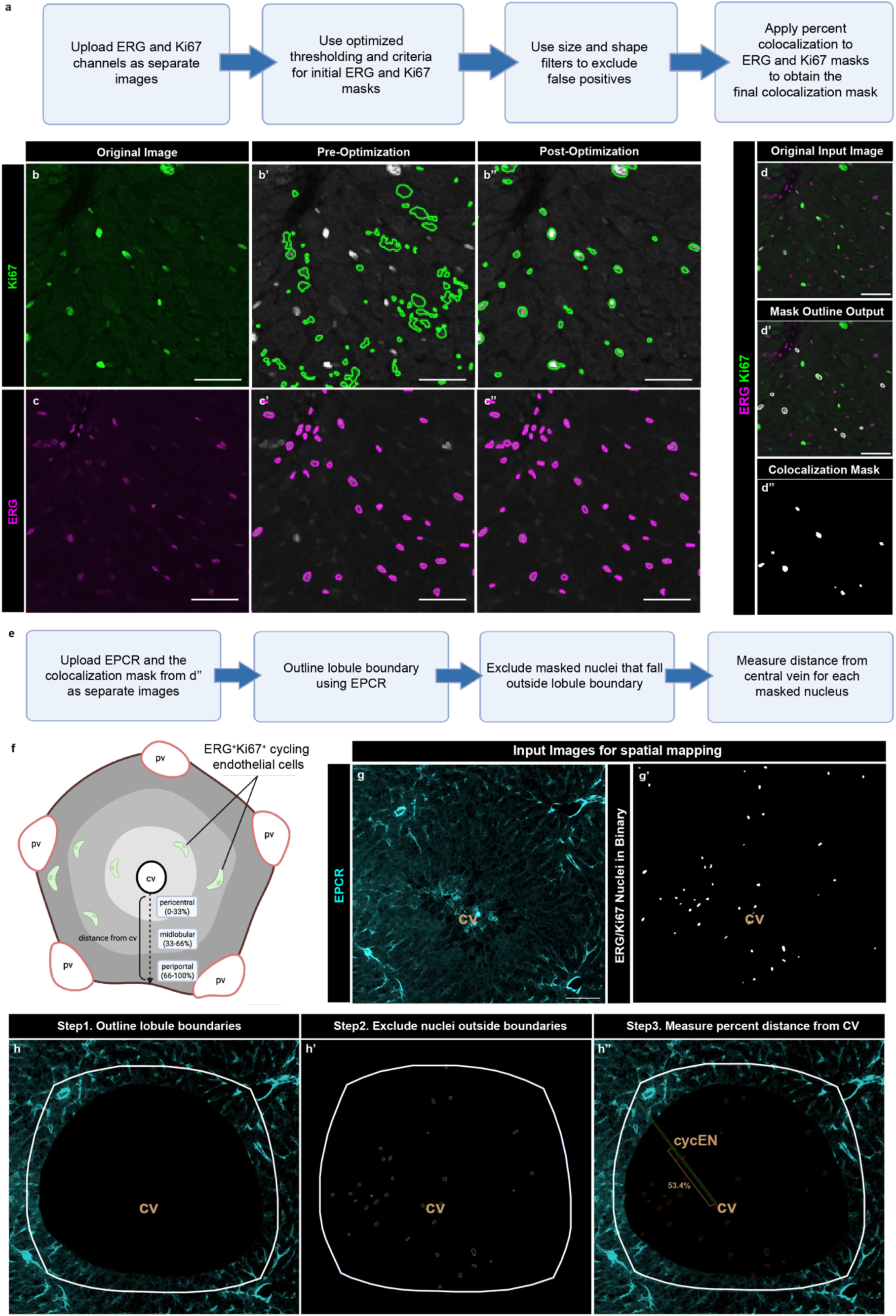
Novel image analysis pipeline enables accurate spatial quantification of cycling cells in liver lobules. **a)** Summary of the first phase of image analysis pipeline where CellProfiler is used to identify ERG^+^/Ki67^+^ nuclei in liver section images. **b-c’’)** Workflow optimization process for identification of Ki67^+^ nuclei in (b-b’’) and for ERG ^+^ nuclei in (c-c’’). Pre-optimization outputs used the default thresholds in CellProfiler. Post-optimization outputs used Ki67 and ERG-specific thresholding method and filters, respectively, as detailed in Methods. Scale bars, 50 µm. (**d-d’’)** Comparison of the input ERG and Ki67 merged image to the mask outline output and the final binary mask output. Scale bars, 50 µm. **e)** Summary of the second phase of image analysis pipeline where MATLAB is used to spatially map the masked nuclei from the first phase. **f)** A schematic summary describing how the zone of individual ERG+/Ki67+ nuclei is calculated in the lobule. The distance along the central-to-boundary axis was divided into three equal parts. Nuclei falling within 0-33 % distance from the central vein are denoted as pericentral, 34-66 % as midlobular, and 67-100 % as periportal. **g-g’)** Input images for the spatial mapping workflow. For each lobule, the Endothelial Protein C Receptor (EPCR) immunofluorescence image and the final binary mask output from (d’’) are fed into the MATLAB workflow. Scale bar, 100 µm. **h-h’)** The steps of MATLAB workflow applied to images in (f-f’). The MATLAB workflow first outlines the lobule boundaries based on EPCR signal. Next, the workflow identifies and excludes masked nuclei that fall outside the lobule boundaries. Lastly, the workflow maps each masked nucleus at a percent distance from the central vein on the central-to-boundary axis. CycEN, cycling endothelial nucleus; CV, central vein; PV, portal vein.

For spatial mapping, we defined the zones within the lobule by calculating percent distance of the nucleus to the central vein on a central-to-boundary axis. The percent distance is used to categorize cells into three zones; pericentral (0-33%), midlobular (34-66%), and periportal (67-100%) (Figure 1f). We chose geometrical categorization over the use of zonal markers to avoid confounding by expression changes, enabling reproducible mapping across injury or disease states. To perform spatial analysis, the ERG^+^Ki67^+^ colocalization output is fed into an in-house generated MATLAB script (Figure 1g-g’). The MATLAB script accurately maps each ERG^+^Ki67^+^ double positive nucleus within the lobule (Figure 1h-h’). This process is applied to as many lobules, biological samples, and experimental conditions as desired in an automated manner. The culminating data provides a rigorous and robust spatial map of cycling cells in a cell-type specific manner in hundreds of lobules across mice. While our study focuses on proliferative response in acute chemical injury, the presented pipeline can be readily adapted to determine the spatial features of cellular or molecular markers of interest in liver lobules in healthy, injured, or diseased states.

### Endothelial cell cycling increases predominantly in the pericentral zone following acute centrilobular injury

Previous work suggested that liver sinusoidal endothelial cells cycle at a higher frequency following acute CCl_4_ injury^22^, focusing on the Lyve1^+^ endothelium that preferentially represents the midlobular zone^26^. A systematic spatiotemporal analysis of pan-endothelial cycling that encompasses the heterogeneity of the lobule vasculature has been lacking. We asked when and where in the lobule endothelial cells enter the cell cycle in response to CCl_4_ by performing pan-endothelial analysis. We harvested livers from injured and vehicle-injected control male mice at progressive time points following treatment and employed the image analysis pipeline from Figure 1. The temporal analysis revealed a rapid peak in endothelial cycling on day 3 following injury (Figure 2a-b). Endothelial cycling remained heightened on day 5 and returned to baseline by day 7 (Figure 2a-b). Prior to the peak, endothelial cycling trended lower than homeostatic levels (Figure 2a-b). These findings suggest that liver endothelial cells at first remain quiescent in response to acute chemical liver injury and afterward rapidly ramp up cell cycle entry.

**Figure 2.**
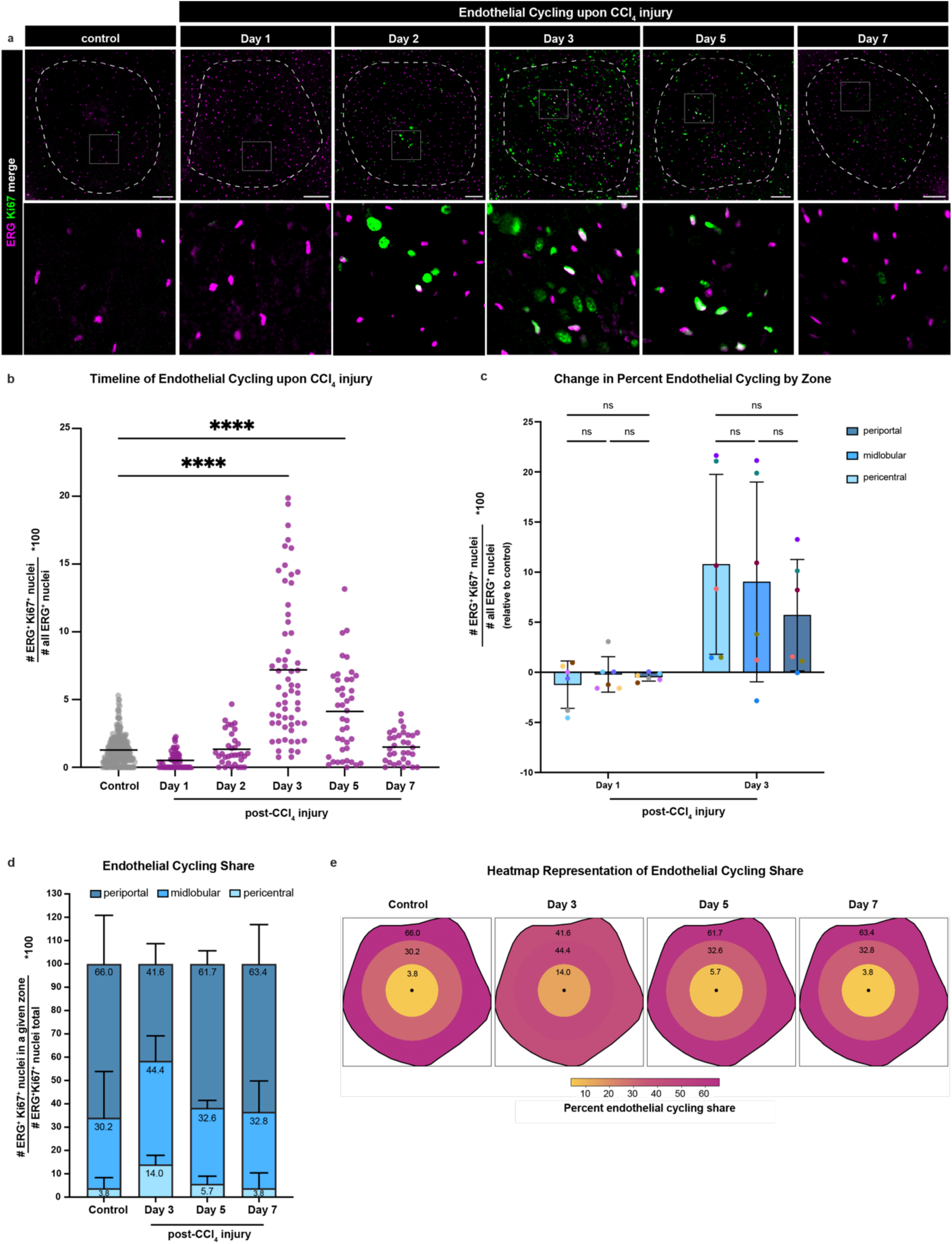
Endothelial cells enter the cell cycle following CCl_4_ injury primarily in the pericentral and midlobular zones. **a)** Immunofluorescence staining of ERG and Ki67 in representative control (corn oil vehicle treated, 1 day post-treatment) and injured livers at progressive timepoints following CCl_4_ injury. Dotted lines delineate lobule boundaries based on EPCR signal. EPCR immunofluorescence not shown for clarity. The lower panels show higher magnification of the corresponding upper panel images. 2 µm thick single slice images are shown. Scale bars, 100 µm. **b)** Quantification of the percent ERG^+^Ki67^+^ nuclei in all ERG^+^ plotted per lobule (n = 265 control lobules, n = 30-60 injury lobules per timepoint pooled from n = 3-5 male mice collected in 3 experimental runs per timepoint). One-way ANOVA with Dunnett’s multiple comparisons test, outliers removed using ROUT method (Q = 1%, 21 datapoints removed) **** p –value <0.0001 **c)** Quantification of change in percent ERG^+^Ki67^+^ double positive nuclei out of total ERG^+^ nuclei within each zone. Change in percent was calculated by subtracting the average value of the control dataset for each zone and plotted by color-coded animals (n = 3-5 animals collected in 3 experimental runs per timepoint). Two-way ANOVA with Šídák’s multiple comparison test was used. **d)** Quantification of percent ERG^+^Ki67^+^ nuclei in each zone out of all ERG^+^Ki67^+^ nuclei in the corresponding lobule, plotted per animal (n = 3-5 collected in 3 experimental runs per timepoint). **e)** Heatmap representing endothelial cycling share over the proliferative period from (d). All graphs show mean +/- SD when applicable. All experiments presented used litter-matched control mice and controls were pooled for analyses comparing multiple time points.

We next asked where in the lobule cycling endothelial cells localize. We performed two parallel spatial analyses. First, we assessed *percent cycling* defined as the percentage of cycling endothelial cells out of all endothelial cells. The percent cycling assessment thus accounts for the difference in the total number of endothelial cells in different zones. All three zones exhibited a similar trend in percent cycling, first declining at day 1 and later increasing at day 3 (Figure 2c). The change in percent endothelial cycling exhibited a very large spread due to interindividual variability, particularly on day 3 (Figure 2c). These data indicate that endothelial cells enter the cell cycle in all zones of the lobule following injury; however, endothelial cycling response varies greatly between individual animals.

In the parallel spatial analysis, we asked how the *cycling share* shifts across zones over time. Cycling share is defined as the percentage of cycling endothelial cells found within a given zone out of total cycling endothelial cells in the entire lobule. This represents the relative contribution of the zones to endothelial cycling, regardless of how many endothelial cells are in each zone. This analysis controls for interindividual variability in the extent of cycling response. At peak cycling at day 3, the endothelial cycling share shifted markedly to the pericentral and midlobular zones from the periportal zone (Figure 2d-e). Midlobular endothelial cells showed the largest cycling share with 44.4 %. When compared to homeostasis, the pericentral zone underwent the biggest change in endothelial cycling share following injury, increasing more than threefold (Figure 2d). The cycling share gradually returned to homeostatic patterns by the end of the proliferative period (Figure 2d-e). These results suggest that the midlobular zone makes the biggest contribution to endothelial cycling following injury, while the pericentral zone undergoes the biggest change in endothelial cycling.

Together, these results indicate that endothelial cycling is altered upon injury most prominently in the pericentral zone, the primary zone of hepatocyte necrosis.

### Hepatocyte cell cycle entry upon centrilobular injury is spatiotemporally distinct from endothelial cell cycle entry

We next asked how endothelial cell cycling correlates with hepatocyte cycling spatially across the lobule and temporally over the regenerative time. Hepatocytes are well-established to rapidly enter the cell cycle following acute chemical liver injury^5,6^; yet, the spatial bias and predominant zone(s) of hepatocyte cycling and lineage expansion continue to be a subject of debate. We leveraged our image analysis pipeline to spatiotemporally map cycling hepatocytes (HNF4α^+^) in molecularly defined lobules upon CCl_4_-injury. The pipeline was optimized to identify the nuclei with Ki67^+^/HNF4α^+^ colocalization through minor threshold and filter adjustments (see Methods for details). Using the optimized pipeline, we first conducted a temporal analysis of hepatocyte cycling. This analysis showed that hepatocytes enter the cell cycle as early as day 2 following injury, in line with previous work^16,19^ (Figure 3a-b). Hepatocyte cycling peaked at day 3 and declined afterwards (Figure 3a-b). Our findings suggest that hepatocytes initiate cell cycle entry earlier than endothelial cells; endothelial cells then rapidly ramp up and peak in cycling concurrently with hepatocytes.

**Figure 3.**
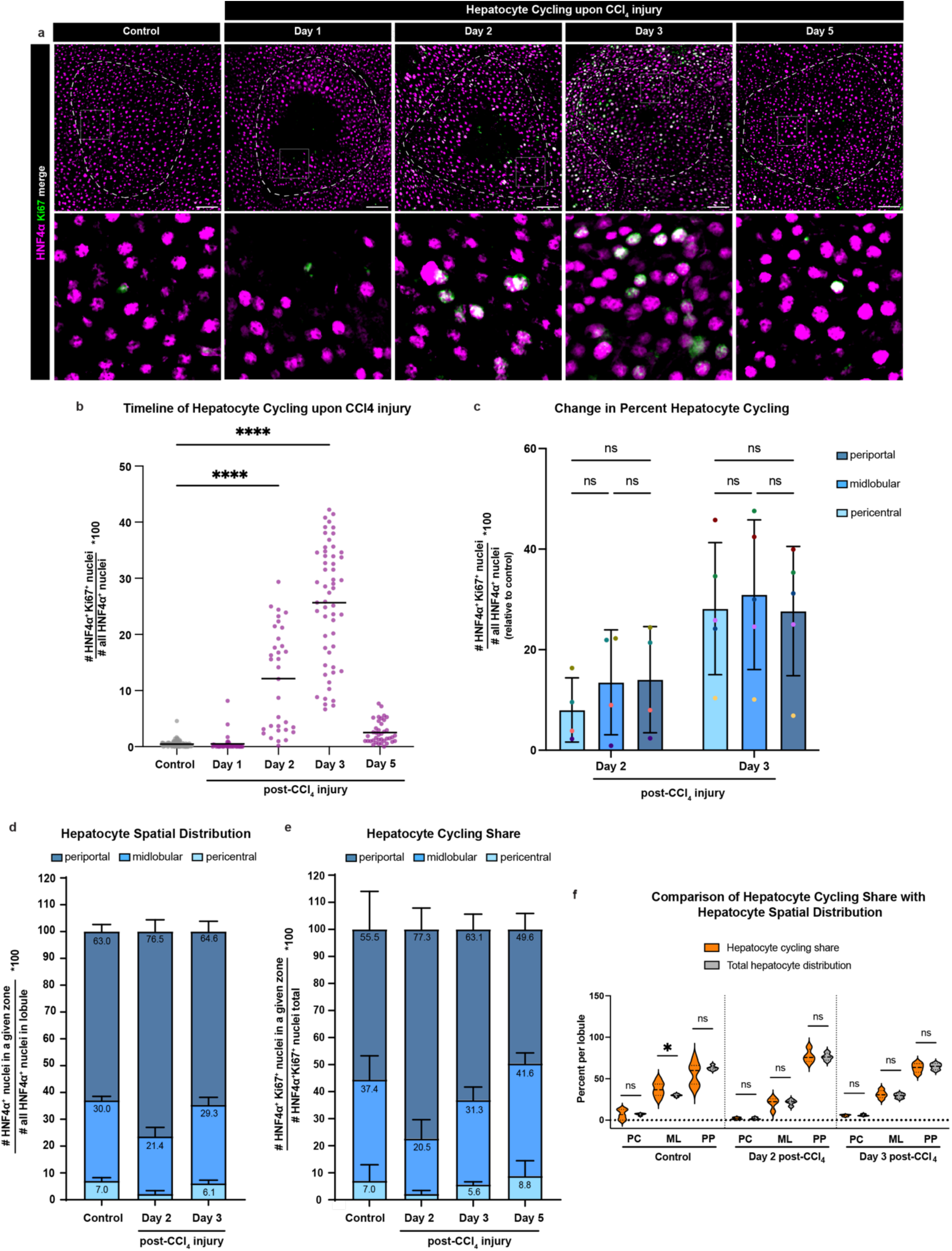
Hepatocytes enter the cell cycle following CCl_4_ injury in a spatiotemporal manner distinct from endothelial cells. **a)** Immunofluorescence staining of HNF4α and Ki67 in representative control (corn oil vehicle treated, 5 days post-treatment) and injured livers at timepoints following CCl_4_ injury. Dotted lines delineate lobule boundaries based on EPCR signal. EPCR immunofluorescence not shown for clarity. The lower panels show higher magnification of the corresponding upper panel images. 2 µm thick single slice images are shown. Scale bars, 100 µm. **b)** Quantification of the percent HNF4α^+^Ki67^+^ nuclei in all HNF4α positive nuclei plotted per lobule (n = 101 control lobules, n = 39-55 injury lobules per timepoint pooled from n = 4-5 male mice collected in 2-3 experimental runs). One way ANOVA followed by Dunnett’s multiple comparisons test was used for statistical analysis. **** p –value <0.0001 **c)** Quantification of change in percent HNF4α^+^Ki67^+^ double positive nuclei out of total HNF4α ^+^ nuclei within each zone, calculated by subtracting the average value of the control dataset for each zone, plotted by color-coded animals (n = 4-5 animals collected in 2-3 experimental runs per timepoint). Two-way ANOVA with Šídák’s multiple comparison test was used. **d)** Quantification of percent HNF4α^+^ nuclei in each zone out of all HNF4α^+^ nuclei in the corresponding lobule, plotted per animal (n = 6 control, n = 4-5 per injury timepoint collected in 2-3 experimental runs). The pericentral average value for day 2 is 2.2%. **e)** Quantification of percent HNF4α^+^Ki67^+^ nuclei in each zone out of all HNF4α^+^Ki67^+^ nuclei, plotted per animal (n = 8 control, n = 4-5 per injury timepoint collected in 2-3 experimental runs). The pericentral average value for day 2 is 2.4%. **f)** Comparison between hepatocyte spatial distribution in (d) and hepatocyte cycling share in (e) by zone; statistical analysis by Kolmogorov-Smirnov test. All graphs show mean +/-SD when applicable. All experiments presented used litter-matched control mice and controls were pooled for analyses comparing multiple time points.

We next investigated how the cycling hepatocytes are distributed across the lobule relative to the cycling endothelial cells. Percent cycling analysis for hepatocytes showed that no single zone has distinctly higher percent cycling than others at either day 2 or day 3 (Figure 3c). While the pericentral zone lagged slightly behind, percent cycling increased in tandem in all three zones from the onset to the peak (Figure 3c). As this analysis relies on the number of all hepatocytes in each zone, we next asked how injury alters the hepatocyte distribution across zones. In injured lobules, periportal hepatocytes constituted most of the remaining hepatocytes (76.4 %), while pericentral hepatocytes were almost completely absent and the midlobular hepatocytes contributed measurably less relative to homeostasis (Figure 3d), as expected from centrilobular damage. Taking these data into account, the apparent increase in percent cycling in the pericentral and midlobular zones results at least in part from lesser number of hepatocytes found in those zones in injured lobules.

We next analyzed hepatocyte cycling share across zones over time (Figure 3e). This analysis revealed that the periportal zone has an inverse trend compared to the other two zones: Cycling share in the periportal zone rapidly increased at the onset of cycling at day 2, then gradually declined (Figure 3e). By contrast, cycling share in the pericentral and midlobular zones declined at first, then gradually increased (Figure 3e). The periportal zone made the biggest contribution to hepatocyte cycling from its onset to its peak (Figure 3e). To determine whether this represents genuine spatially biased cycling, we asked how cycling share compares to the spatial distribution of hepatocytes. We reasoned that, if cycling share diverged from the distribution of total hepatocytes, this would suggest genuine spatially biased cycling. By contrast, if cycling share tracks with the distribution of total hepatocytes, this would suggest that cycling is not spatially biased; rather, the abundance of hepatocytes in each zone simply dictates the abundance of cycling hepatocytes. Distribution analyses yielded no quantifiable difference between hepatocyte cycling share and total hepatocytes in injury, neither at the onset nor at the peak of hepatocyte cycling (Figure 3f). These results suggest that hepatocyte cycling in centrilobular injury predominantly occurs in the periportal zone, not due to genuine spatially biased cycling, rather because this zone contains majority of the remaining healthy hepatocytes following centrilobular injury.

We conclude that hepatocyte cycling does not exhibit genuine spatial bias in any zone in centrilobular injury; rather, it occurs uniformly across healthy hepatocytes remaining in the lobule. Taken together with our results on endothelial cycling predominantly heightened in pericentral and midlobular zones, these findings indicate that hepatocyte cycling is spatially uncoupled from endothelial cell cycling in acute chemical liver injury.

### Endothelial cells divide and expand over the regeneration period following acute centrilobular injury

Cell cycle entry has been reported to occur without complete division in response to injury in other tissues^27^. As endothelial cells are not thought to be the primary target of hepatotoxin injury^28,29^, we hypothesized that endothelial cell cycle entry upon acute CCl4 injury occurs in the absence of complete division or cellular expansion. We tested whether cycling endothelial cells go on to divide and expand in injury using the pan-endothelial Ve-Cadherin^CreERT2^ in combination with the sparse labelling reporter Confetti, referred to as VE-Cad^CreERT2^;Confetti (Figure 4a). Utilizing volumetric imaging, we performed three-dimensional pan-endothelial clonal expansion analysis. Contrary to our hypothesis, this analysis revealed that a higher number of endothelial cells divide and expand in injured livers compared to controls (Figure 4b-c). In injured livers, 50 % more endothelial cells underwent cell division (Figure 4c). The frequency of endothelial cells that divided once, resulting in two-cell clones, was similar between injury and control (Figure 4d). By contrast, the endothelial cells that formed three- or more-cell clones were almost exclusive to injured livers. These results demonstrate that endothelial cells clonally expand in response to acute chemical liver injury and identify serial endothelial cell division as a distinguishing characteristic of injured livers.

**Figure 4.**
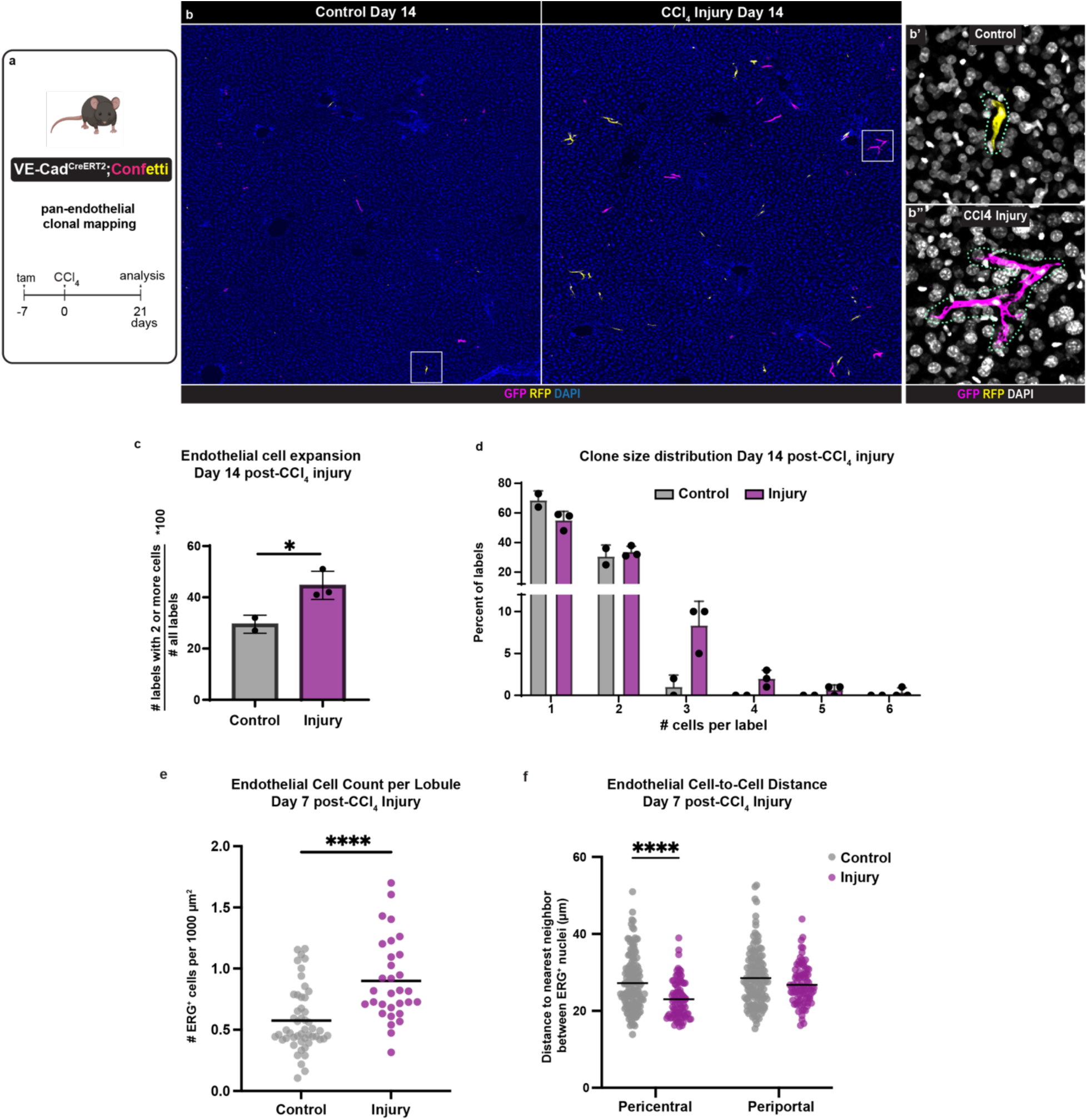
Endothelial cells divide and expand during liver regeneration from CCl4 injury. **a)** Experimental scheme of pan-endothelial clonal mapping experiments upon CCl_4_ injury using VE-Cadherin^CreERT2^;Confetti mice. **b)** Immunofluorescence staining of GFP and RFP to mark the lineage-labelled endothelial cells 14 days after CCl_4_ injury or control (corn oil vehicle) treatment. DAPI marks the nuclei. 32 µm thick maximum intensity projection images are shown. **c)** Quantification of percentage of labels expanded 14 days after CCl_4_ injury or control treatment plotted per animal. Two-tailed unpaired t test was used for statistical analysis. * p –value <0.05. **d)** Distribution of clone sizes determined by the number of cells per label, plotted per animal. **e)** Quantification of total ERG^+^ cells in a lobule 7 days after CCl_4_ injury, plotted per lobule (n = 50 control lobules, n = 31 injury lobules, pooled from n = 3-5 male mice collected in 3 experimental runs). Two-tailed unpaired t-test was used for statistical analysis. **** p –value <0.0001. **f)** Quantification of nearest neighbor distance between ERG^+^ nuclei in pericentral and periportal areas 7 days after CCl_4_ injury. Each dot represents a region of 100 µm ^2^ in the pericentral or periportal zone (n = 287 control, n = 177 injury, pooled from n = 3-5 male mice collected in 3 experimental runs). Two-way ANOVA with Sidak’s multiple comparisons test between control versus injury and pericentral versus periportal zones. **** p –value <0.0001. All graphs show mean +/-SD when applicable.

To determine how endothelial cell expansion affects lobule composition, we asked whether lobules of injured livers are altered in endothelial cell count or density. To this end, we quantitatively assessed total ERG^+^ cell count at the end of the proliferative period upon injury. This analysis identified a significant increase in endothelial cell count in lobules of injured livers compared to controls (Figure 4e). In parallel, we analyzed endothelial cell density by measuring nucleus-to-nucleus distance within the endothelium. Indeed, endothelial nuclei were more densely packed in injured liver lobules, more pronounced in the pericentral zone (Figure 4f).

Our findings altogether suggest that endothelial cell cycle entry predominantly localized to the necrotic zone followed by endothelial clonal expansion results in overabundance of endothelial cells in regenerating lobules upon acute centrilobular liver injury.

## Discussion

The liver vasculature and its specialized endothelial cells are paramount to liver structure and function. Previous work has shown that liver sinusoidal endothelial cells respond to acute chemical liver injury by entering the cell cycle^22^. However, due to the lack of definitive lobule markers and of reproducible spatial quantification methods, it has been challenging to identify which zonal endothelial populations contribute to liver recovery. In this study, we develop a novel automated image analysis pipeline that accurately maps cell cycling in liver lobules and perform systematic characterization of the proliferative endothelial response to acute centrilobular liver injury.

Previous spatial mapping in the liver has widely relied on hepatocyte gene expression for zonal definitions, most commonly Glutamine Synthetase for the pericentral zone and E-Cadherin for the periportal zone^12,13,15,16,19^. These studies have provided important insights into the spatial relationship between hepatic gene expression, cell states, and blood vessels. Yet, genuine zonal information in the context of the native lobule structure has been challenging to attain. This can be attributed to a combination of factors: First, hepatocyte gene expression is dynamically altered following injury^17,20,30^, and does not consistently provide a reliable point of reference. Second, hepatocyte gene expression along the central-portal axis is not binary but gradual^31^, confounding reproducibility. Third, transverse sectioning widely used in previous work does not account for the native hexagonal lobule architecture^23,24^, causing sampling artifacts and misrepresentation of the relative zonal contributions. The image analysis pipeline we present circumvents these challenges. The presented pipeline utilizes a definitive marker of lobule boundaries on coronal liver sections and geometrically maps cells within the native architecture of hexagonal lobules. The automated high-throughput nature of the presented pipeline helps generate reproducible and robust spatial maps of cell types and states in liver lobules.

Our studies demonstrate that endothelial cells enter the cell cycle upon acute centrilobular injury at a later time than hepatocytes, in line with previous findings on sinusoidal endothelial cycling^22^. Spatially, our results indicate that endothelial cells enter the cell cycle most dynamically in the pericentral zone followed by the midlobular zone, the zones of parenchymal damage. A previous transcriptomics study found a cycling profile in the pericentral endothelium upon acetaminophen exposure^17^, similar to our findings. These results indicate that the biggest change in endothelial cycling occurs near parenchymal damage, supporting the notion that endothelial cycling may occur in response to hepatocyte necrosis.

Further studies are needed to directly test this and alternative hypotheses. Our findings additionally present the first line of evidence, to our knowledge, that endothelial cells go on to divide and clonally expand upon parenchymal liver injury. Serial cell division emerges as a distinguishing behavior of endothelial cells of injured livers, rarely seen in homeostasis. Unexpectedly, prior to cell cycle entry, endothelial cells trend towards a decline from homeostatic cycling at first, suggesting that a subset of endothelial cells might exit the cell cycle rapidly following parenchymal injury. The molecular triggers and mechanisms underlying endothelial cell cycle exit, entry, and clonal expansion in acute chemical liver injury remain to be defined by future work.

Our work suggests that hepatocyte cell cycle entry upon acute centrilobular injury occurs uniformly across three zones of the liver lobule. Previously, multiple studies identified slight or substantial spatial bias in hepatocyte proliferation upon centrilobular injury^16,17,19–21^, while others proposed that all zones contribute uniformly to hepatocyte proliferation^13^ in line with the findings presented here. Our study expands on previous work and demonstrates the importance of considering total hepatocyte numbers in each zone when assessing genuine spatial bias, particularly in injury models. In addition, our results indicate enriched hepatocyte proliferation in the midlobular zone in the homeostatic liver and thus support previous lineage-tracing studies^13,15,16,19^. Interestingly, both hepatocyte and endothelial proliferative responses exhibit high interindividual variability in the absence of a litter effect. Further analyses are needed to determine whether the variability in proliferative response is caused by a corresponding variability in the extent of initial damage. These results highlight the need to consider interindividual variability in the analysis of proliferative response to liver injury. Our findings prompt further investigation into the zonal contributions to hepatocyte and endothelial proliferative response in other types of chemical, physical, and chronic injury.

Overall, our study establishes a novel automated image analysis pipeline to spatiotemporally map cell states in liver lobules, and identifies a zonally biased proliferative response in the endothelium to acute centrilobular liver injury. These findings contribute to a better understanding of the multifaceted endothelial response to acute liver injury. Future work on the endothelial biology of acutely injured livers will provide insights into the mechanisms underlying endothelial adaptation in contrast with maladaptation, and facilitate a holistic approach towards the treatment of chronic liver injury patients.

## Methods

### Mice

The Institutional Animal Care and Use Committee at University of North Carolina at Chapel Hill approved all animal procedures and protocols, conformed to the Guide for the Care and Use of Laboratory Animals. Experiments were performed with adult male 8 to 12-week-old mice. Weaning age was set at 23-25 days. All mice were housed under a 12 h:12 h light:dark cycle at 22–25 °C and 30–70% humidity. Mice received rodent chow and water ad libitum. C56BL/6 mice were used for wild-type studies. Acute liver injury was induced by intraperitoneal injection of CCl_4_ at a dose of 1mL/kg body weight (1:4 dilution in corn oil). Corn oil was injected intraperitoneally using the same volume by weight calculation as a vehicle control. Confetti (017492) mice were purchased from The Jackson Laboratory and bred in-house. VE-CadherinCreERT2, also called Cdh5-PAC-CreERT2 or Cdh5-cre/ERT21Rha, mice were generated by the Adams Lab^32^. CreERT2 mice were crossed into Confetti mice for clonal mapping studies.

### Tamoxifen treatment and lineage tracing studies

Adult male VE-Cadherin^CreERT2^;Confetti mice were induced with tamoxifen by intraperitoneal injection at 8-12 weeks of age. Tamoxifen was dissolved in corn oil with 10% ethanol at a concentration of 5mg/mL and administered a single dose of 1mL/kg.

### Vibrating microtome sectioning and immunostaining

Thick section processing and immunostaining protocol was adapted from Snippert et al, 2011. For embedding and sectioning, Precisionary compresstome equipment and tools were used (VF-310-0Z).

Livers were harvested and fixed in 4% paraformaldehyde overnight at 4 °C followed by PBS washes (3 × 5 min). The whole left median lobe was embedded curvature down. The embedding was performed in the manufacturer’s mold in 4% ultrapure low-melting agarose prepared in PBS. The samples were left in the embedding mold for 10 min followed by incubation in the provided ice block for 5 min. The samples were sectioned at 100 µm at speed set to 3 and vibration set to 4.

For immunostaining, compresstome sections were permeabilized in PBS containing Triton X-100 0.1% for 1 h, blocked in PBS with Triton X-100 0.1% containing normal donkey serum at 5% for 1 h, and incubated in primary antibodies for 2 overnights at 4 °C. The sections were then washed in PBS containing Triton X-100 0.1% (4 x 15 min) and incubated in secondary antibodies raised in donkey conjugated to Alexa Fluor 488, 555, or 647 (1:400; Jackson Immunoresearch Labs) overnight at 4 °C. Next, the sections were washed in PBS containing Triton X-100 0.1% (4 x 15 min), incubated in DAPI solution in PBS (1:1000, ThermoFisher, D1306) for 30 min, and washed again for 15 min. Lastly, using a paintbrush, the sections were mounted on a slide with Prolong Gold with DAPI (Invitrogen) and protected with a coverslip.

For HNF4α and ERG antibodies, heat-mediated antigen retrieval was performed following permeabilization and prior to blocking, as previously described^24^. Antigen retrieval was performed in Tris Buffer (Vector Labs H-3301) in a pressure cooker (for ERG) or oven (for HNF4α). Following cool-down, sections were blocked. After this step, the sections were incubated in primary antibodies, and the regular compresstome staining protocol was followed.

### Confocal imaging

Immunostained compresstome liver sections were imaged using a Leica SP8 confocal. 20X multi-immersion objective with glycerol and 40X oil objective were used. Images of full liver sections were tiled and stitched by automatic merge using the LasX Software (4.9.2). Z stacks of images were projected into a 2-dimensional image when applicable by applying maximum intensity projection in FIJI (ImageJ).

### Antibodies

The following primary antibodies were used at indicated concentrations for the immunostaining of compresstome sections: goat anti-EPCR (R&D Systems, AF2749, 1:100), mouse anti-Hepatocyte nuclear factor 4 alpha (Abcam, ab41898, 1:200), chicken anti-GFP (Abcam, ab13970, 1:500), rabbit anti-RFP (Rockland, 600-401–379, 1:200), and rabbit anti-ERG (Abcam, ab110639, 1:200), rat anti-Ki67 (Invitrogen,14-5698-82, 1:500).

### Image processing, analysis, and quantification

Tiled and stitched full-section confocal images of z-stacks (32 um) were converted into maximum intensity projection images, and the projected images were used for further analyses unless otherwise noted below. Whole left median lobes were used in all experiments unless otherwise noted below or in the figure legends. Same thickness images were used across samples for quantification. For the presentation and quantification of ERG immunostaining, the images were despeckled in FIJI (ImageJ). Image quantifications are described in detail below.

### Colocalization analysis of ERG^+^Ki67^+^ and HNF4α^+^Ki67^+^ nuclei

Single z-slice images from EPCR, Ki67 and ERG or HNF4α-immunostained full liver sections were cropped into images of individual lobules identified using EPCR for colocalization analysis. n = 265 control lobules (pooled for all timepoints) and n = 30-60 injury lobules per timepoint from 3-5 male mice collected in 3 experimental runs were analyzed in Figure 2. n = 101 control lobules (pooled for all timepoints) and n = 39-55 injury lobules per timepoint from 4-5 male mice collected in 2-3 experimental runs were analyzed in Figure 3. The custom CellProfiler workflows generated in this study for colocalization analysis of endothelial cells and hepatocytes are publicly available and can be accessed at https://github.com/azizoglulab/Liver_Injury_Cell_Cycling.

### Spatial distribution of ERG^+^Ki67^+^ and HNF4α^+^Ki67^+^ nuclei

Binary outputs from colocalization analysis of ERG^+^Ki67^+^ or HNF4α ^+^Ki67^+^ were used. The binary mask and EPCR channel were run through a custom MatLab script generated in this study. n = 17 control mice (pooled for all timepoints) and n = 3-4 injury mice per timepoint collected from 3 experimental runs were analyzed in Figure 2. n = 8 control mice (pooled for all timepoints) and n = 4-5 injury mice per timepoint collected from 2-3 experimental runs were analyzed in Figure 3. The custom MatLab script generated in this study for spatial distribution analysis is publicly available and can be accessed at https://github.com/azizoglulab/Liver_Injury_Cell_Cycling.

### Spatial distribution of ERG^+^ and HNF4α^+^ nuclei

Cropped individual lobule images of single z-slice were run through the same custom CellProfiler workflow used for colocalization analysis above which by default calculates the number of ERG+ or HNF4α+ nuclei. The binary outputs were run through the same custom MatLab script as described for the spatial distribution of ERG+Ki67+ and HNF4α+Ki67+ nuclei above.

### Quantification of ERG^+^ nuclei count

Cropped individual lobule images of single z-slice were used. For each image, ERG channels were run through the same custom CellProfiler workflow as above. The obtained number of ERG^+^ nuclei was then normalized to the area of the corresponding cropped image. N = 50 control lobules and n = 31 injury lobules per timepoint from 3-5 male mice collected in 3 experimental runs were analyzed in Figure 4.

### Quantification of endothelial nucleus-to-nucleus distance

ERG, EPCR, and Ki67-immunostained full liver section single z-slice images were cropped into three 100 µm^2^ squares per pericentral region for 10 central veins identified. For each image, ERG channel was separated and run through a custom CellProfiler workflow generated in this study to calculate the form factor of ERG^+^ cells. n = 287 control and n = 177 injury cropped regions from 3-5 male mice collected in 3 experimental runs were analyzed in Figure 4. The custom CellProfiler workflow generated in this study for nucleus-to-nucleus distance analysis of endothelial cells is publicly available and can be accessed at https://github.com/azizoglulab/Liver_Injury_Cell_Cycling.

### Quantification of Vecad^CreERT2^;Confetti clonal expansion

GFP, RFP, and EPCR-immunostained 32 µm-thick full liver sections were used for analysis. GFP and RFP labelled clones were marked and numbered. A random number generator was utilized to randomly select 100 clones per animal for further analysis. The labelled clones were quantified for the number of cells contained within each clone using the whole 32 µm z-stack. n = 2 and n = 3 injury male mice collected in 3 experimental runs were analyzed.

### Statistical information

Statistical analyses were performed using Prism 10. Statistical methods and sample sizes are detailed in the corresponding figure legends.

## Code Availability

The custom Matlab script and CellProfiler pipelines have been deposited to Github and can be accessed at https://github.com/azizoglulab/Liver_Injury_Cell_Cycling (10.5281/zenodo.23771487).

## Acknowledgements

The authors thank the members of the Azizoglu lab for their input and feedback. This manuscript was funded by the American Liver Foundation through the ALF Liver Scholar Award.

## Author Contribution Statement

ASR and HY performed most of the experiments and analyses. EJH and EPS performed endothelial proliferation studies and analyses with supervision from ASR and HY. ASR, HY, and KLT prepared the original draft and reviewed and edited the manuscript with input from DBA. DBA conceptualized, acquired funding, supervised, and validated the studies and conclusions.

